# Evaluating three different adaptive decomposition methods for EEG signal seizure detection and classification

**DOI:** 10.1101/691055

**Authors:** Vinícius R. Carvalho, Márcio F.D. Moraes, Antônio P. Braga, Eduardo M.A.M. Mendes

## Abstract

Signal processing and machine learning methods are valuable tools in epilepsy research, potentially assisting in diagnosis, seizure detection, prediction and real-time event detection during long term monitoring. Recent approaches involve the decomposition of these signals in different modes or functions in a data-dependent and adaptive way. These approaches may provide advantages over commonly used Fourier based methods due to their ability to work with nonlinear and non-stationary data. In this work, three adaptive decomposition methods (Empirical Mode Decomposition, Empirical Wavelet Transform and Variational Mode Decomposition) are evaluated for the classification of normal, ictal and inter-ictal EEG signals using a freely available database. We provide a previously unavailable common methodology for comparing the performance of these methods for EEG seizure detection, with the use of the same classifiers, parameters and spectral and time domain features. It is shown that the outcomes using the three methods are quite similar, with maximum accuracies of 97.5% for Empirical Mode Decomposition, 96.7% for Empirical Wavelet Transform and 98.2% for Variational Mode Decomposition. Features were also extracted from the original non-decomposed signals, yielding inferior, but still fairly accurate (95.3%) results. The evaluated decomposition methods are promising approaches for seizure detection, but their use should be judiciously analysed, especially in situations that require real-time processing and computational power is an issue. An additional methodological contribution of this work is the development of two python packages, already available at the PyPI repository: One for the Empirical Wavelet Transform (ewtpy) and another for Variational Mode Decomposition (vmdpy).

## Introduction

Epilepsy is a burdening neurological disease that has a prevalence rate of around 6 per 1000 persons and incidence rate of 61 per 1000 person-years [1]. One of the factors contributing for its high incidence rate is the large number of causes leading to this condition, such as: genetic predisposition, displasias, cerebrovascular disease (CVD), trauma, tumor, infection, ischemia, among others [2].

Recurrent seizures are considered the hallmark of Epilepsy. These events reflect the abnormal firing of groups of neurons in the brain, in general synchronous and of high intensity [3]. This deviation from the normal functioning patterns of neurons may invoke sensations varying from strange feelings, behaviors and sensations to seizures with muscular spasms and possible loss of conscience [4].

The electroencephalogram (EEG) is a high temporal resolution recording of brain electrical activity central to the diagnosis of epilepsy and other neurological disorders. Its signals can reflect abnormal neuronal activity during ictal (i.e. seizures) or interictal periods, such as sharp transients occurring in-between seizures [5]. These signals are commonly interpreted by experienced neurologists through visual inspection, taking into account features such as frequency, amplitude and regularity of waveforms, reactivity to stimuli, spatial range and temporal persistence of the signal’s anomalies [6]. However, this method may be cumbersome and time consuming, especially for long series and multi-channel data; which can lead to an increasingly high ratio of false-negative results. Furthermore, there is a series of subtle signal features and components, as well as inter-channel relationships, which are virtually impossible to detect by simple visual inspection. This task may be assisted by signal processing and classification algorithms [7] that can deal with signal nonlinearities and subtleties, high-dimensional data and the possibility of real-time processing. As such, these automated methods are valuable tools for the diagnosis, detection and prediction of epilepsy and epileptic seizures [8].

A variety of algorithms and signal processing techniques have been developed for the extraction of relevant features related to the epileptic phenomena [9]. Methods which analyze frequency components using the Fourier Transform are not always recommended, because EEG signals contain non-stationary components, violating conditions for the use of such transform [10]. Thus, recent methods for EEG analysis of epileptic patients may use Time-Frequency approaches, or non-linear methods such as Lyapunov Exponents, Fractal Dimension, Entropy or Correlation Dimension [11]. Other methods include the use of signal decomposition in adaptive ways, such as the Empirical Mode Decomposition (EMD), proposed by Huang et al [10].

Of particular interest to the work presented here, the EMD is an adaptive and data-dependent decomposition method that successively extracts intrinsic mode functions (IMFs), defined by amplitude modulated (AM) and frequency modulated (FM) components. Accordingly, complex non-linear and non-stationary signals can be decomposed into a finite number of IMFs, each with well-behaved Hilbert Transforms [10]. The EMD approach, as well as its extensions [12,13], has been successfully used in epilepsy [14–17]. However, drawbacks such as computational cost, lack of theory (due to its algorithmic approach) and difficulty in interpreting the large number of modes have motivated the use and evaluation of different adaptive decomposition methods in seizure EEG signals [18].

The Empirical Wavelet Transform (EWT) [18] addresses some limitations of EMD. By adapting some of the Wavelet formalisms, this method designs appropriate wavelet filter banks and decomposes a signal into a predetermined number of modes. The use of EWT has been explored in different areas such as compression of electrocardiogram (ECG) signals [19], decomposition of seismic activity [20] and time-frequency representation of non-stationary signals [21]. Although the use of Wavelets for seizure detection and classification has been widely explored [6,22–24], few works evaluate EWT for processing seizure EEG signals [25,26].

Another adaptive method denominated Variational Mode Decomposition [27] (VMD) decomposes a signal into its principal modes adaptively and non-recursively. The method is related to the so-called Wiener filter, a property which grants it advantages in relation to noise robustness. Similar to what happened in the case of EWT, few researchers have evaluated VMD use for seizure EEG analysis [28–30].

When processing EEG from ictal phenomena, features generated from the use of the aforementioned decomposition methods are promising tools. In addition to their adaptive capabilities and ability to deal with nonlinear and non-stationary signals, extracted modes (or signal components) are compact around specific center frequencies and have well-behaved Hilbert transforms. This enables the extraction of features related to amplitude or bandwidth modulation, as well as instantaneous phase and amplitude. This work aims to compare these three decomposition methods for EEG signal seizure detection using a freely available database and a common methodology, by extracting the same features and using the same classifiers. So far, this comparison has been hampered not only by the small number of works using EWT and VMD, but also by the fact that, for seizure detection, EMD, VMD and EWT are evaluated in the literature using different sets of features, parameters and classifiers. In this work, results are also compared with features extracted from original non-decomposed signals. Expected results are performance improvements with the use of adaptive decomposition methods (EMD, EWT and VMD), but similar overall performances among them.

The rest of this work is organized into three sections. In Section 2, the used methodology is presented, containing the used data, decomposition methods, description of extracted features and classification problem. In Section 3, the obtained results are presented and discussed. Concluding remarks are left for the last section.

## Methods

### Dataset

The EEG data used in this work was obtained from a public online database offered by the University of Bonn. This dataset is divided into 5 subsets: *Z, O, N, F, S* (or A, B, C, D, E) [31]. Each subset contains 100 temporal series with sampling frequency of 173.6 Hz and duration of 23.6 seconds. The Z and O subsets correspond to surface EGG recordings of 5 healthy volunteers, with eyes open and closed, respectively. The rest of the subsets belong to presurgical recordings of epileptic patients. Set S contains seizure activity, while sets F and N are from seizure-free intervals, with electrodes placed on the epileptogenic zone and opposite hippocampus, respectively.

This work deals with the most common classification problem, using three subsets: S, F and N, corresponding to Ictal, Interictal and Normal classes, respectively. Samples from each class and their regularized spectra, given by the Gaussian-filtered Fast Fourier Transform (FFT) of each signal are shown in **Erro! Fonte de referência não encontrada.**.

**Fig 1.**
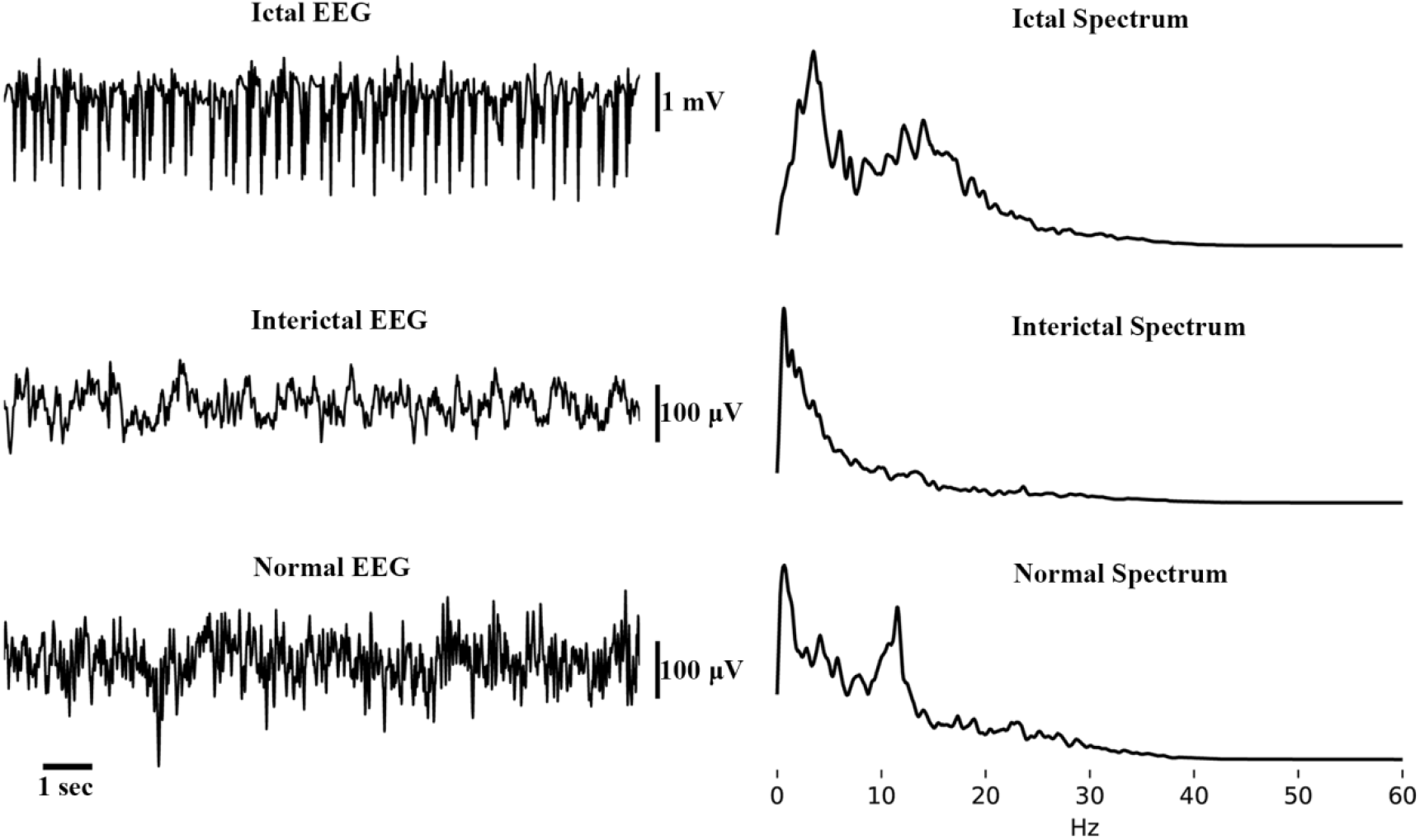
Example EEG signals of each class (Ictal, Interictal and Normal) and respective spectra

The first stage of processing consists of applying a zero-phase 4^th^ order lowpass Butterworth filter, with cutoff frequency equal to 40 Hz. Next, signals are decomposed into N modes or IMFs, either by EMD, EWT or VMD, which are described in the following sections.

### Empirical Mode Decomposition

The Empirical Mode Decomposition (EMD) is an interesting method due to its adaptability, not depending on assumptions as linearity or stationarity. This method aims to divide the analyzed signal into a series of Intrinsic Mode Functions (IMFs), where each IMF must satisfy to two relatively simple conditions:

I. The number of extrema must be equal or differ by at one (at most) in relation to the number of zero crossings.
II. In every sample, the mean envelope value, defined by the local maxima and minima, must be equal to zero.

The algorithm proposed by [10] for obtaining IMFs consists of the following steps:

1. Given a temporal series *x*(*t*), find the local maxima and minima.
2. Generate the envelopes *e*_*max*_ and *e*_*min*_ by cubic spline interpolation of maxima and minima, respectively.
3. Calculate the mean of the envelopes,*m*_*i*_(*t*) =(*e*_*max*_(*t*) + *e*_*min*_(*t*))/2
4. Subtract the value found previously from the: *h*(*t*) = *x*(*t*) − *m*_*i*_(*t*) If *h*(*t*) satisfies the conditions given previously for an IMF, an IMF *c*_*i*_(*t*) = *h*(*t*) is found.
5. A new residue *r* is generated: *r*(*t*) = *x*(*t*) − *c*_*i*_(*t*) Repeat steps 1 to 4, applied to the residue *r*, in order to find the remaining IMFs. The process halts when it is no longer possible to compute an IMF from a residue, which is then defined as a final residue *r*_*M*_.

The signal is then decomposed into a determined number of IMFs *c*_*i*_(*t*), plus another residue *r*_*M*_, and represented by (1),

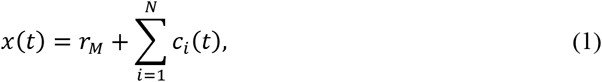

where N is the total number of IMFs found.

Unlike methods such as the Discrete Wavelet Transform, which extracts low frequencies (or approximations) first, and detail levels (corresponding to higher frequencies) later, the first modes isolated by EMD correspond to high frequencies of the signal, then moving progressively to slower components.

The pyEMD Python package [32] was used for the implementation of EMD in this work.

### Empirical Wavelet Transform

As in EMD, EWT method aims to extract the oscillatory amplitude (AM) and frequency (FM) components of a signal, considering these as having compact Fourier support. Unlike traditional wavelet transforms, which use predefined filter bank structures, EWT defines the supports of the filters in accordance with the spectral distribution of the signal, in a fully adaptive way. Some considerations are made for analysis: (1) the signal must be real valued, due to the need for symmetry, and (2) a normalized frequency axis with 2π periodicity is considered, but analysis is restricted to [0, π], due to Shannon’s sampling criterion.

A number of modes N is defined a priori, determining how many segments the spectrum is partitioned in the range [0, π]. Among the N+1 frequency limits to be determined, two are already predefined (*ω*_*0*_ and *ω*_*N*_), corresponding to frequencies of 0 and π, respectively. The remaining N-1 limits are set according to the distribution of the signal’s frequency spectrum: the N-1 local maxima are found, and the limits *ω*_*n*_ (n = 1,2, .. N-1) are defined as midpoints between two consecutive maxima. In this work, maxima were detected on the smoothed spectrum that is obtained by applying a Gaussian filter (filter length = 25, σ = 5) on the Fast Fourier Transform of each signal. The segmentation of a signal spectrum into 5 Modes is given in **Erro! Fonte de referência não encontrada..**

**Fig 2.**
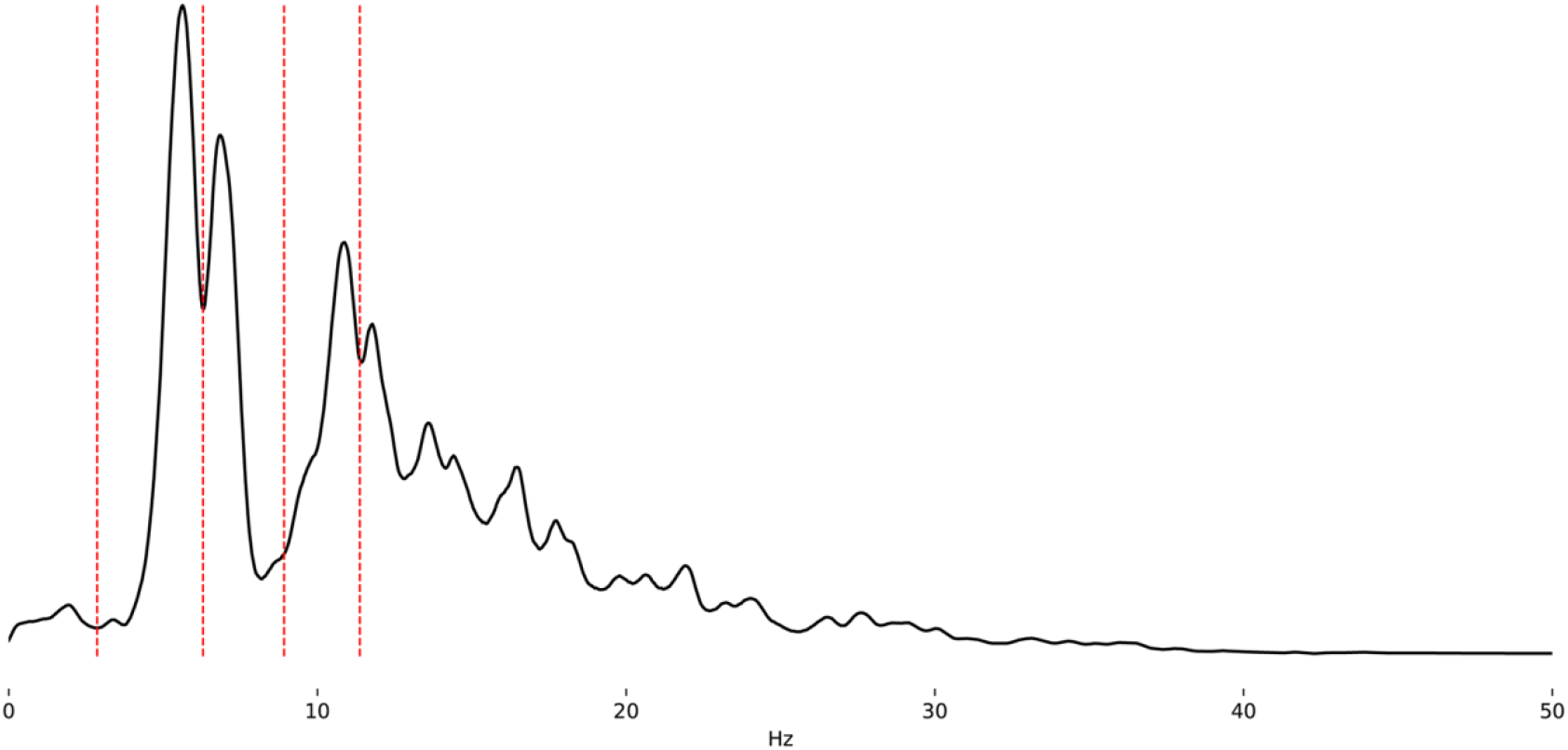
Spectral segmentation of ictal EEG into 5 modes

With limits *ω*_*n*_ defined, the segments *Λ*_*n*_ = [*ω*_*n-1*_, *ω*_*n*_] fill the interval [0, π]. The limits of each segment are characterized by a transition period centered at the respective *ω*_*n*_, with width equal to 2τ_n._ Each segment is associated to a filter (lowpass for the *ω*_*0*_, bandpass for the rest), the construction of which is related to Littlewood-Paley and Meyer Wavelets [33]. Thus, an empirical scale function *φ*_*n*_(*ω*) and an Empirical Wavelet *ψ*_*n*_(*ω*) are defined. These are constructed in such a way that a Tight Frame is obtained. Further details on the construction of such functions are given in [18].

With the conditions for building a Tight Frame satisfied, the EWT is defined similarly as the traditional Wavelet Transform, with details given by the inner product of the Wavelet function with the signal, and the approximation equal to the inner product of the signal with the scaling function.

Based on Jérôme Gilles’ MATLAB toolbox [34], a Python package of EWT (ewtpy) was developed for this work and is available at https://pypi.org/project/ewtpy/ and at https://github.com/vrcarva/ewtpy.

### Variational mode decomposition

In the Variational mode decomposition, the number of modes is predefined. Initially, the method assumes each mode *k* as having compact bandwidth in the Fourier spectrum, with a respective central frequency. *ω*_*k*_ For each mode, a unilateral frequency spectrum is obtained and shifted to baseband according to its estimated central frequency. The bandwidth is then assessed by the H^1^-norm Gaussian smoothness of the demodulated signal, with the optimization problem iteratively updating each mode in the frequency domain. The complete constrained variational optimization problem is available in [27].

In comparison with EMD, VMD performed better in tests dealing with tone detection and separation, and noise robustness [27]. And although its use for long-time EEG signals suffers from caveats due to non-stationarity, the use of VMD in this work is motivated by the relatively short duration of the EEG signals from the used Database, and the focus on classification rather than exact mode decomposition and reconstruction.

A Python package, based on the original MATLAB toolbox [35], was developed for this work and made available at https://pypi.org/project/vmdpy/ and at https://github.com/vrcarva/vmdpy. These can be readily installed with pip (*pip install vmdpy* and *pip install ewtpy*).

### Hilbert Transform and Analytic signal

The presented decomposition methods have the interesting property of yielding modes/IMFs with “well-behaved” Hilbert transforms [10,27], from which features as instantaneous phase, frequency or envelope can be extracted. The analytic signal of each mode/IMF is given by equation (2).

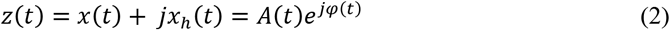

where *x*_*h*_(*t*) is the Hilbert transform of *x*(*t*),

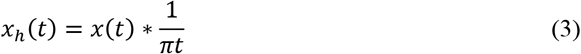

and the amplitude and phase are defined as:

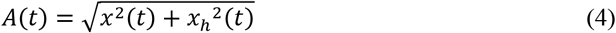

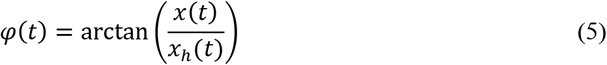

The instantaneous frequency of a given IMF can be calculated from its analytic signal with:

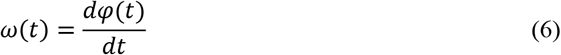

By dividing a signal into a given number of modes/IMFs, features related to their respective analytic signals and spectra can be extracted.

### Feature extraction

Modes given by the three aforementioned methods (EMD, EWT and VMD) may be considered as amplitude and frequency modulated signals. Thus, feature extraction is made according to properties of the spectrum of each mode, with a similar approach used by [36] and [15].

The first feature extracted is the Spectral Energy (SE), given by Equation 7.

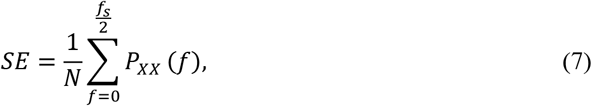

where N is the total number of spectral coefficients, and *P*_*XX*_ is the mode PSD estimated by Welch’s method [37]. The second feature is the Spectral Entropy (SEnt), shown in Equation 8.

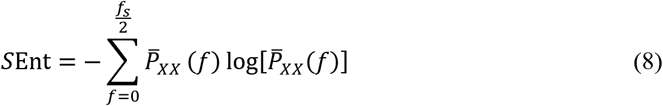

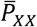 is the normalized PSD. The following three features are related by the main frequency component of the respective mode. After determining the global maximum of *P*_*XX*_, the corresponding magnitude is defined as the Spectral Peak (SP), as well as the associated frequency (*f*), defining the 3^rd^ and 4^th^ features. The following feature is the spectral centroid (SC) of the respective mode, defined in Equation 9:

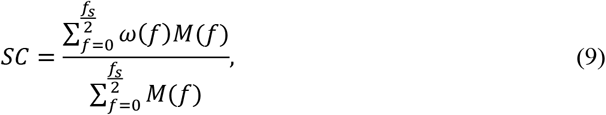

where *f* is the frequency bin, and *ω*(*f*) and *M*(*f*) are respectively, the central frequency and magnitude of the PSD of bin *f*. The last two features are the AM and FM bandwidths, defined by Equations 10 and 11 [38]:

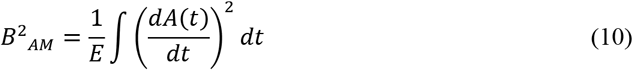

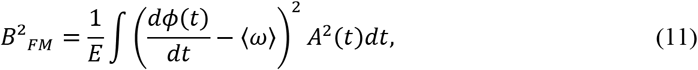

where *A* is the amplitude of the analytic signal, *E* is the Energy and 〈*ω*〉 is the center frequency of the current mode, given by Equation 12.

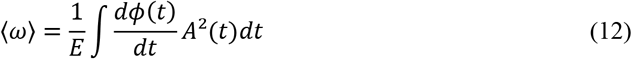

Time-domain features are also extracted from each mode: Hjorth parameters [39] and statistical moments.

Hjorth Mobility is related to the mean frequency of the signal and proportional to the variance of its spectrum, while Hjorth Complexity is an estimate of the signals’ bandwidth [40]. These are defined by:

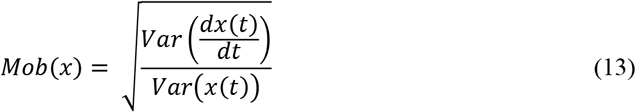

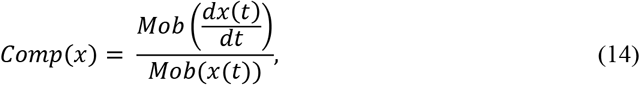

where *x*(*t*) is the current signal component, and Var() is the variance.

The Skewness is related to the signal distribution’s asymmetry, and is given by the following equation:

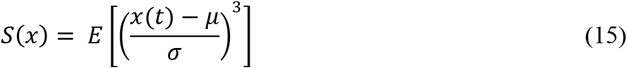

The standard deviation of *x*(*t*) is represented by *σ*, and its mean by *μ*. The Kurtosis is related to the tails of the distribution yielded by the signal and is given by.

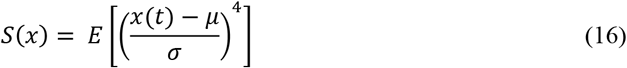

### Feature selection and classification

For feature selection and classification algorithms, functions from scikit-learn package [41] were used.

Since the number of extracted features of each signal is relatively large, and not every feature is relevant for class discrimination, there is a need for a feature selection or ranking method. In this work, 15 features were selected according to the SVM Recursive Feature Elimination (RFE) [42]. Afterwards, different classification methods were evaluated: k-nearest neighbors (KNN) [43], Linear and Radial Basis Function (RBF) Support Vector Machines (SVM) [44], Gaussian process classification (GPC) based on Laplace approximation [45] and a Multi-layer Perceptron (MLP) [46].

In order to avoid data overfitting, 5-fold cross-validation was used for the classification algorithms, with 70% of samples used for Training, and 30% for testing. For performance evaluation, four common measures were employed; Accuracy (ACC), Specificity (SPEC), Sensitivity (SEN) and Area Under the Receiver Operating Characteristic (ROC) curve (AUC). The last three measures are calculated by choosing the ictal class as positive.

## Results and discussion

All scripts used in this work are available online at https://github.com/vrcarva/carvalho-etal-2019.

Fig 3 illustrates the decomposition of a seizure EEG by EMD, EWT and VMD. Fig S1, and Fig S2 show this for inter-ictal and normal signals, respectively.

**Fig 3.**
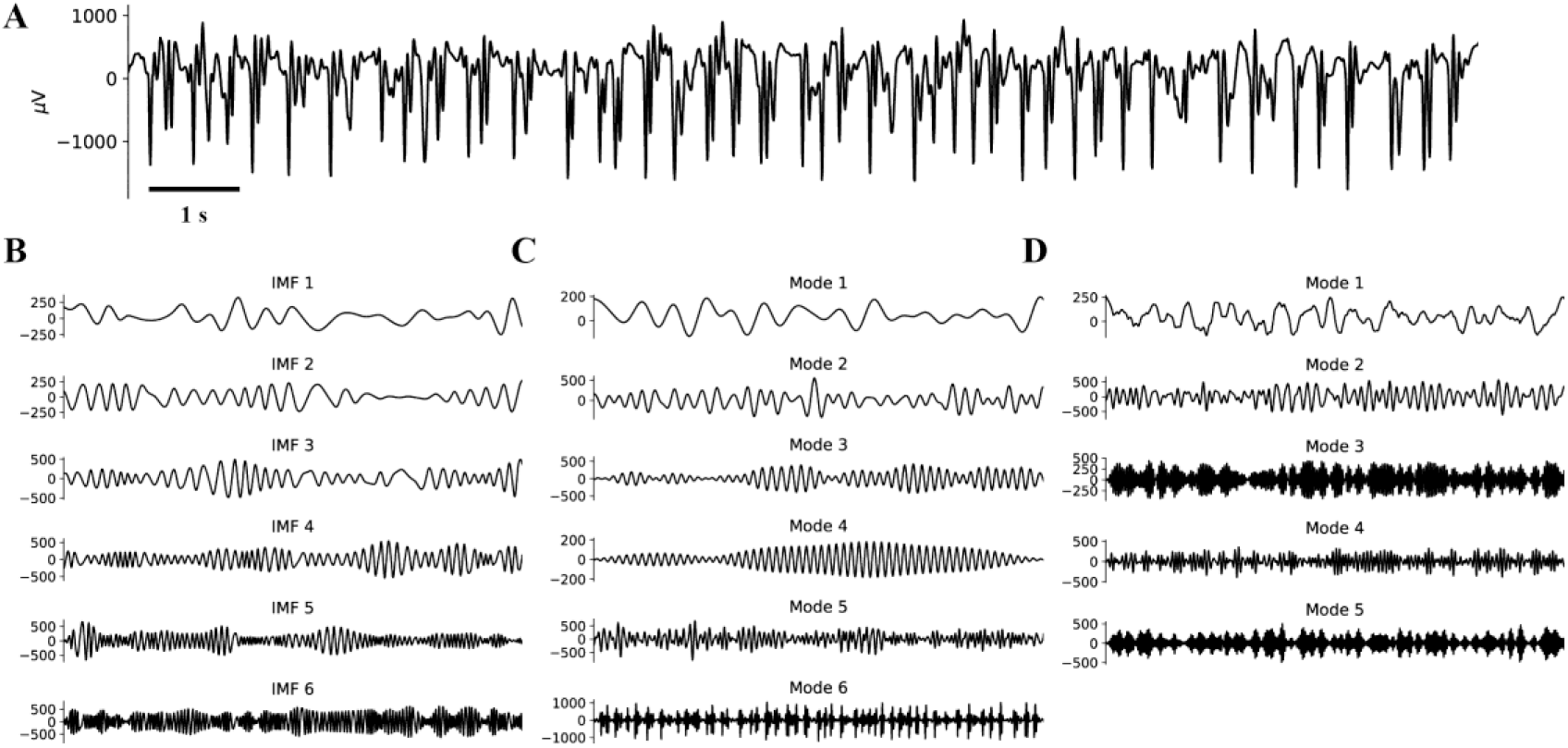
Decomposition of ictal EEG. (A) Original non-decomposed signal. (B) First six extracted IMFs with EMD. (C) 6 components extracted with EWT. (D) 5 components extracted with VMD.

Samples of each class were decomposed by EMD, EWT or VMD, into N (4, 5, 6 and 7) Modes, followed by the extraction of the 11 features of each one, thus resulting in N*11 features for each sample. The SVM-RFE algorithm then selected 15 of these features. RFE was not applied for the “control” (the non-decomposed signal), which results in 11 features for each sample. Afterwards, different classifiers are trained and tested, resulting in performance evaluation metrics. Ten iterations of training/testing were accomplished, resulting in mean ± standard deviation for each performance parameter. The best results for each classifier applied to each decomposition method are shown in Table 1.

**Table 1:** Classification results of different decomposition (EMD, EWT, VMD and original signal) and classification methods.

| METHOD | MODES | MEASURE | KNN | LINEAR SVM | RBF SVM | GPC | MLP |
| --- | --- | --- | --- | --- | --- | --- | --- |
| EMD | 6 | AUC (%) | 99.00±0.28 | 99.25±0.22 | 99.63±0.11 | 99.72±0.08 | 99.55±0.14 |
|  |  | ACC (%) | 97.10±0.63 | 95.90±0.33 | 97.53±0.22 | 97.0±0.58 | 96.80±0.34 |
|  |  | SEN (%) | 94.20±1.60 | 94.10±0.83 | 98.60±0.49 | 96.2±0.98 | 95.10±0.94 |
|  |  | SPEC (%) | 99.05±0.35 | 97.30±0.24 | 97.50±0.22 | 97.9±0.58 | 98.15±0.23 |
| EWT | 6 | AUC (%) | 96.97±0.61 | 99.27±0.08 | 99.35±0.09 | 99.32±0.17 | 99.43±0.10 |
|  |  | ACC (%) | 93.80±0.93 | 95.77±0.42 | 96.60±0.70 | 95.93±0.51 | 96.70±0.38 |
|  |  | SEN (%) | 92.60±1.36 | 94.50±1.02 | 96.00±0.63 | 95.6±1.20 | 95.80±1.08 |
|  |  | SPEC (%) | 94.40±1.04 | 96.45±0.35 | 97.20±0.40 | 96.55±0.47 | 97.60±0.44 |
| VMD | 5 | AUC (%) | 98.17±0.31 | 99.25±0.18 | 99.27±0.11 | 99.49±0.19 | 99.41±0.10 |
|  |  | ACC (%) | 96.83±0.37 | 96.90±0.40 | 98.20±0.31 | 97.50±0.50 | 97.67±0.42 |
|  |  | SEN (%) | 95.90±1.30 | 95.80±0.87 | 97.40±0.49 | 96.60±0.80 | 95.70±0.64 |
|  |  | SPEC (%) | 97.70±0.46 | 97.85±0.45 | 98.65±0.39 | 98.45±0.57 | 98.85±0.23 |
| - | 1 | AUC (%) | 97.31±0.51 | 96.93±0.31 | 98.24±0.29 | 98.36±0.31 | 98.12±0.28 |
|  |  | ACC (%) | 93.27±0.79 | 89.87±0.65 | 92.60±0.42 | 95.27±0.73 | 93.27±0.51 |
|  |  | SEN (%) | 91.10±1.76 | 89.00±0.63 | 88.70±1.55 | 93.60±1.20 | 89.70±0.90 |
|  |  | SPEC (%) | 94.85±1.10 | 90.80±0.81 | 95.05±0.52 | 96.10±0.66 | 95.50±0.84 |

Best-case results are similar among all three decomposition methods, with slight superior values for EMD and VMD. RBF SVM yielded the best performance, with ACC of 97.5%, 96.7% and 98.2% for EMD, EWT and VMD, respectively. These results are in accordance with the literature, which has accuracy values varying from 90% in [47] to 99,7% in [48]. VMD and autoregression (AR) based quadratic feature extraction with a random forest classifier was used in [29], resulting in up to 96.4% ACC. Random forests with VMD were also used in [28], that combined with semantic feature extraction resulted in a maximum accuracy of 94.1%. Entropy measures in [49] achieved 92.8% and 91.0% ACC. Broader summaries of different methods and respective performance measures are given in [50] and in [36]. The latter uses EMD and similar features of the ones used in this work, achieving 95.7% ACC, 98% SEN and 97% SPEC.

The methods presented in this work aim to decompose a signal into AM-FM components, and extract features of each one, which may differentiate “normal” signals from pathologic ones. This has promising applications for the identification of epileptogenic systems and for identifying interictal-ictal state transitions. The feature extraction capabilities can be further explored if each mode was considered as having “well behaved” Hilbert Transforms, which enables the analysis of phase relationships between different modes of the same signal or between different brain regions, if multichannel data is available. Since synchronization is believed to play an important role for ictogenesis, these methods may further assist on unveiling the mechanisms of seizure generation and devising markers to predict transitions to ictal states.

Although the results are positive, the application of the presented methods for epileptic seizure detection and classification must face a series of challenges. The Bonn University EEG database is useful for preliminary algorithm evaluation and comparison with similar works. However, considering that the data consists of selected short segments obtained in controlled conditions, it has light requirements in terms of generalization and robustness in comparison with what would be found in a clinical environment. Nevertheless, it is an important evaluation step that should be considered for seizure detection and prediction methods.

## Conclusion

The development of feature extraction and classification methods is a key step both for understanding of the operating mechanisms of epilepsy, as for clinical analysis, including possible applications for seizure detection and prediction devices.

This work has shown that EWT and VMD, relatively new signal decomposition methods, may be used to extract relevant features for classifying and detecting EEG signals of epileptic phenomena. Although EMD and VMD presented the best performance results, the trade-off falls on higher computational power, which may compromise its application for real-time processing. The EWT method, in spite of its slightly worse performance, may be preferable if the context of use that requires faster processing.

The use of adaptive decomposition methods is a promising approach due to their ability to separate different AM-FM components that are altered in the presence of seizures (and possibly on periods preceding these events). This would facilitate the extraction of features related to these events, increasing the performance of classification algorithms. However, the extraction of the same features from of non-decomposed signal still results in fairly accurate classifiers. Thus, in contexts with the need of real time processing and limited computational power, the use of these decomposition methods might not be needed for detecting seizures in cases with less strict performance requirements.

Another goal of this work was to provide a Python code of these signal decomposition methods for the community. Python is a fast-growing programming language and is currently the third most popular programming language in the world [51] and with widespread applications in neuroscience [52]. The distribution of these packages could further encourage the use of open source programming languages for works involving these specific signal processing methods.

## Acknowledgements

This work was financially supported by CAPES - Brazil. We also thank CNPq and FAPEMIG for the research support and development.

---

**Fig S1. Decomposition of inter-ictal EEG.** (A) Original non-decomposed signal. (B) First six extracted IMFs with EMD. (C) 6 components extracted with EWT. (D) 5 components extracted with VMD.

**Fig S2. Decomposition of normal EEG.** (A) Original non-decomposed signal. (B) First six extracted IMFs with EMD. (C) 6 components extracted with EWT. (D) 5 components extracted with VMD.

