## Supplementary figures and images for "Evaluating three different adaptive decomposition methods for EEG signal seizure detection and classification"

### Supplemental Figure 1

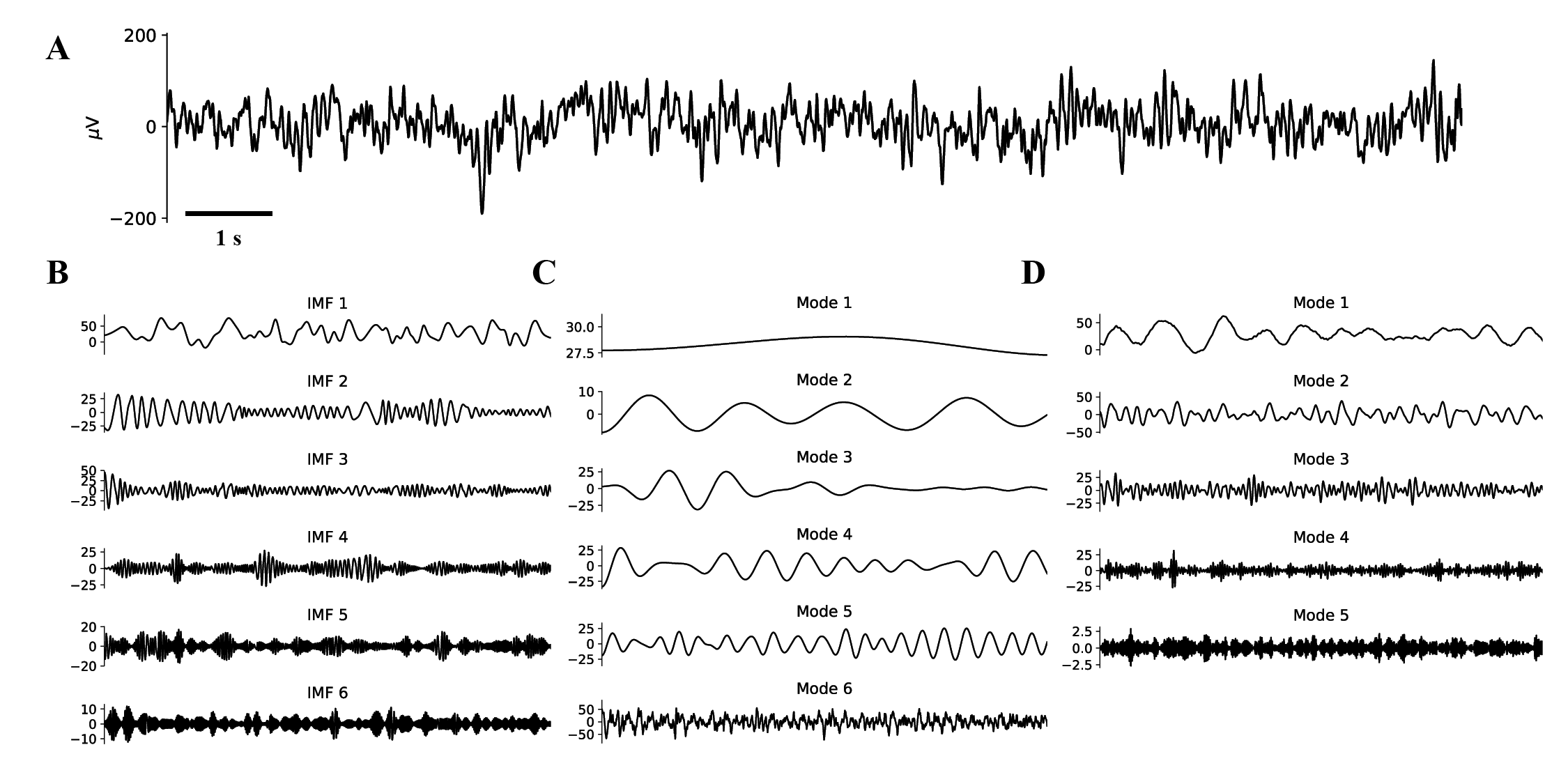

### Supplemental Figure 2

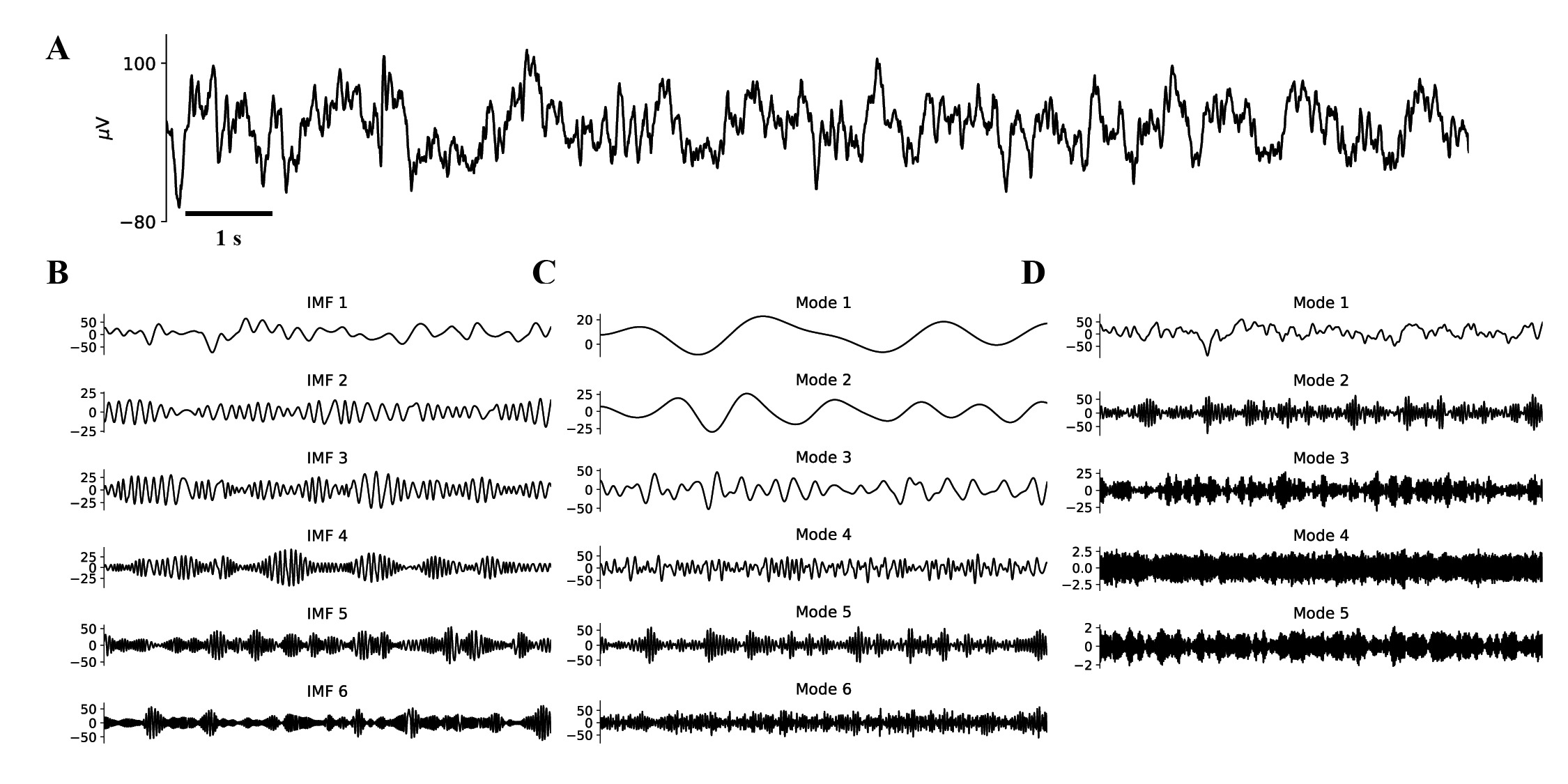
